# Epithelial Tissues as Active Solids: From Nonlinear Contraction Pulses to Rupture Resistance

**DOI:** 10.1101/2020.06.15.153163

**Authors:** Shahaf Armon, Matthew S. Bull, Avraham Moriel, Hillel Aharoni, Manu Prakash

## Abstract

Epithelial tissues in many contexts can be viewed as soft active solids. Their active nature is manifested in the ability of individual cells within the tissue to contract and/or remodel their mechanical properties in response to various conditions. Little is known about the emergent properties of such materials. Specifically, how an individual cellular activity gives rise to collective spatiotemporal patterns is not fully understood. Recently we reported the observation of ultrafast contraction pulses in the dorsal epithelium of *T*.*adhaerens* in vivo [1] and speculated these propagate via mechanical fields. Other accumulating evidence suggest mechanics is involved in similar contractile patterns in embryonic development in vivo and in cellular monolayers in vitro. Here we show that a widespread cellular response – activation of contraction in response to stretch – is sufficient to give rise to nonlinear propagating contraction pulses. Using a minimal numerical model and theoretical considerations we show how such mechanical pulses emerge and propagate, spontaneously or in response to external stretch. The model – whose mathematical structure resembles that of reaction-diffusion systems – explains observed phenomena in *T. adhaerens* (e.g. excitable or spontaneous pulses, pulse interaction) and predicts other phenomena (e.g. symmetric strain profile, “spike trains”). Finally, we show that in response to external tension, such an active two-dimensional sheet lowers and dynamically distributes the strains across its surface, hence facilitating tissue resistance to rupture. Adding a cellular softening-threshold further enhances the tissue resistance to rupture at cell-cell junctions. As cohesion is at the heart of epithelial physiology, our model may be relevant to many other epithelial systems, even if manifested at different time/length scales.

**Significance:** Our work demonstrates that many observed dynamical phenomena in epithelial tissues can be explained merely by mechanical cell-cell interactions, and do not require chemical diffusion or transport between cells (though chemical activity may participate in relevant intracellular processes). Specifically, we show that single cell extension-induced-contraction (EIC) is sufficient to generate propagating contraction pulses, which also increase the tissue’s resistance to rupture, an essential function of epithelia. Our results may shed light on how epithelial tissues function under challenging physiological conditions, e.g. in lung, gut, vasculature and other biomedical contexts. Our results may also be relevant in the study of early evolution of multicellularity and the nervous-muscular systems. Finally, the work offers guidelines for designing soft synthetic solids with improved mechanical properties.

## Introduction

Epithelial cells form confluent layers and are thus inherently mechanically coupled. In recent years multiple observations are accumulating showing contraction patterns and dynamics in epithelial tissues that are suspected to be governed mechanically. These evidence include contraction fronts in drosophila embryo during development [2] and density waves in MDCK cell monolayers in-vitro, either in confinement [3, 4], during expansion [4, 5] or under substrate-shear [6]. Theoretical works suggest models to explain the contractile patterns [7–11], models that include mechanical alongside chemical fields, diffusion or active transport. Note, that all these systems, and the suggested models, exhibit dynamics in time scales of minutes to hours.

Recently we reported the observation of ultrafast contraction dynamics in the dorsal epithelium of *T. adhaerens*, including traveling pulses that propagate across the entire animal [1]. The pulses are seen while the animal is freely moving. The propagating pulses initiate spontaneously, not regularly, propagate at a speed of 1-3 cells per second (10-30um/sec), and transmit a contraction of about 50% in cell area in 1 second (Fig1a-c). These contraction waves can propagate uniaxially, split at the propagating fronts and annihilate themselves. Importantly, this early-divergent animal has no reported muscles, neurons or synapses, and its epithelium has no gap junctions that can support cell-cell transport. In addition, since the propagation speeds are extremely fast, it excludes slower processes (such as transcription) from being involved. All these raise the speculation that mechanics govern the contraction propagation.

*T. adhaerens* is an ideal system to investigate epithelium dynamics. The thin, flat, suspended monolayer resembles early embryonic tissues not only in its minimal components (lacking extra cellular matrix and basement membrane) but also in its large strains dynamics (both individual cells and the entire body undergo dramatic shape changes constantly) [1]. However, unlike embryonic tissues, which undergo slow, stereotypical deformations in time scale of minutes to hours, contraction pulses in the dorsal epithelium of *T. adhaerens* propagate in time scale of seconds, in what seems to be part of a physiological/responsive behavior: due to the animal’s erratic, locally-driven ciliary locomotion, the tissue is found constantly under alternating tensile/compression stresses [1]. In very large animals the *ventral* epithelium, that does not show contraction behavior, does fracture [12].

The main candidate to be involved in such fast contraction dynamics is Calcium. While Calcium is known to immediately activate actomyosin contractions in neurons and muscles, less is known about calcium signaling in epithelia [13]. Nevertheless “calcium waves” have been observed in various epithelial systems, such as drosophila embryo imaginal disc during development [13, 14] and mammalian cornea, in response to mechanical stress or injury [15, 16]. These waves refer to propagation of high cytoplasmic calcium levels from cell to cell via gap junctions. However, the hierarchy between chemical, mechanical and electrical signals is not entirely clear, as mechano-sensitive Calcium channels are known to be involved [15, 17].

In live, behaving *T. adhaerens* we showed [1] that increasing intracellular calcium levels (by adding the drug Ionomycin to the media) yields simultaneous contractions of all cells to ∼ 50% area within seconds [1]. Hence, it is likely that the spontaneous contractions involve calcium as well. However, it is unclear how one contraction triggers the other during a contraction pulse in an early animal that lacks synapses or gap junctions [18]. Diffusion of neuro-peptides in the surrounding water was suggested to coordinate ciliary beating in the ventral tissue of this animal [19], but chemical diffusion is not likely to govern the contraction pulses in the dorsal epithelium, due to the nature of their propagation (uniaxial propagation, front split, pulses annihilation etc).

Here we propose a mechanical model for inter-cellular transmission of contraction in epithelia. The model is based solely on the common single-cell response of Extension-Induced Contraction (EIC). Various experimental evidence have shown such responses in different systems, and different molecular mechanisms have been proposed to participate in it [4, 20–28]. These include mechanosensitive Calcium channels, ERK protein activation, actin alignment, myosin recruitment, conformational changes in adherens junction and more. The diversity of mechanisms for EIC generation hints about its possible evolutionary advantages. We frame our model regardless of the specifics of these intra-cellular processes, and suggest that cellular EIC may be the only requirement for contraction propagation. As such, our model may be relevant to many epithelial systems.

We present numerical and analytical results for tissue contraction dynamics emerging from cellular EIC. Primarily, we write scaling laws for contraction pulses, map its different modes in parameter space, and show that the dynamic patterns are governed by reaction-diffusion type of equations. The model may explain many phenomena seen in *T. adhaerens*, such as single or repeated pulses, either oscillatory or irregular, and pulse annihilation. We utilize the model to predict effects that should be confirmed experimentally, including “spike trains” with fixed pulse amplitude and symmetric pulse profile.

Lastly, a surprising outcome of our model is the emerging enhanced resistance to rupture in such active materials. We show that the contractile dynamics resulting from EIC support tissue cohesion, as it evenly distributes external loads throughout the surface, reducing high strain values. Note, that this homogenization consumes cellular energy. Also note, that the even strain distribution increases the stresses at cell-cell junctions. In addition, it has been shown in various epithelial systems that the cell cytoskeleton responds to mechanical stretch not only by stiffening/contraction (EIC), but also by softening/yielding [28–32]. How do these antagonist trends interplay is still unknown. By numerically introducing a cell softening response to our model (on top of EIC) we further enhance cohesion, by eliminating high stress values at the junctions. Together, the two cellular responses simultaneously prevent tissue failure in a process we name “active cohesion”.

## Results

We begin by looking at the recently-measured contraction profile in *T. adhaerens* [1] (fig1c). The profile shows the strain evolution of a single cell during a single contraction event (as averaged from multiple cellular events that did not propagate as pulses in the tissue). On average, during such a contraction event, cell area increases gradually to a critical point, at which it abruptly decreases to about half its initial value. Finally, the cell relaxes slowly towards its initial area. Despite the overdamped conditions, the contraction reaches significantly below the cell’s steady state size (“overshoot”). This shows that the contraction is an active process (that consumes internal cell energy) and suggests it includes a force and time scales that are different from the viscoelastic ones. As the contraction seem to happen at some critical cell size, an additional scale is required, to describe the extension needed for activation. These three parameters are the minimal requirements for the model.

We therefore model a single cell as an overdamped elastic entity (rest length *l*_0_, elastic modulus *k*, media viscosity *γ*) that is connected in parallel to an active contractile unit (fig1d). The active unit is realizing a simple EIC, using the three scales discussed above: when the cell reaches a critical length *l*_*c*_, it immediately applies a fixed compression force *f*_*c*_ for a duration of *t*_*c*_ (see discussion for further reasoning and alternative options, see methods for implementation). Inspired by *T. adhaerens* data, we take the active forces to be larger than the elastic force at criticality (*f*_*c*_ > *kl*_*c*_) and the active time scale shorter than the passive one (*t*_*c*_ < *k*/*γ*). Finally, we assume the spring to be linearly elastic: effects of cell-volume conservation, though shown irrelevant to the thin, soft tissue of *T. adhaerens* [1], may require additional nonlinearities.

The mechanical circuit proposed in fig 1d can be found in one of two modes: an *excitable mode*, i.e. contract only in response to external stretch (that is when *l*_*c*_ > *l*_0_), or in an *oscillatory mode*, i.e. oscillate spontaneously without any external stimulation, as a relaxation oscillator (that is when *l*_*c*_ < *l*_0_)(fig1e). Such an isolated-cell behavior can be mathematically described in a piecewise manner (fig 1D, SI1).

**Fig. 1:**
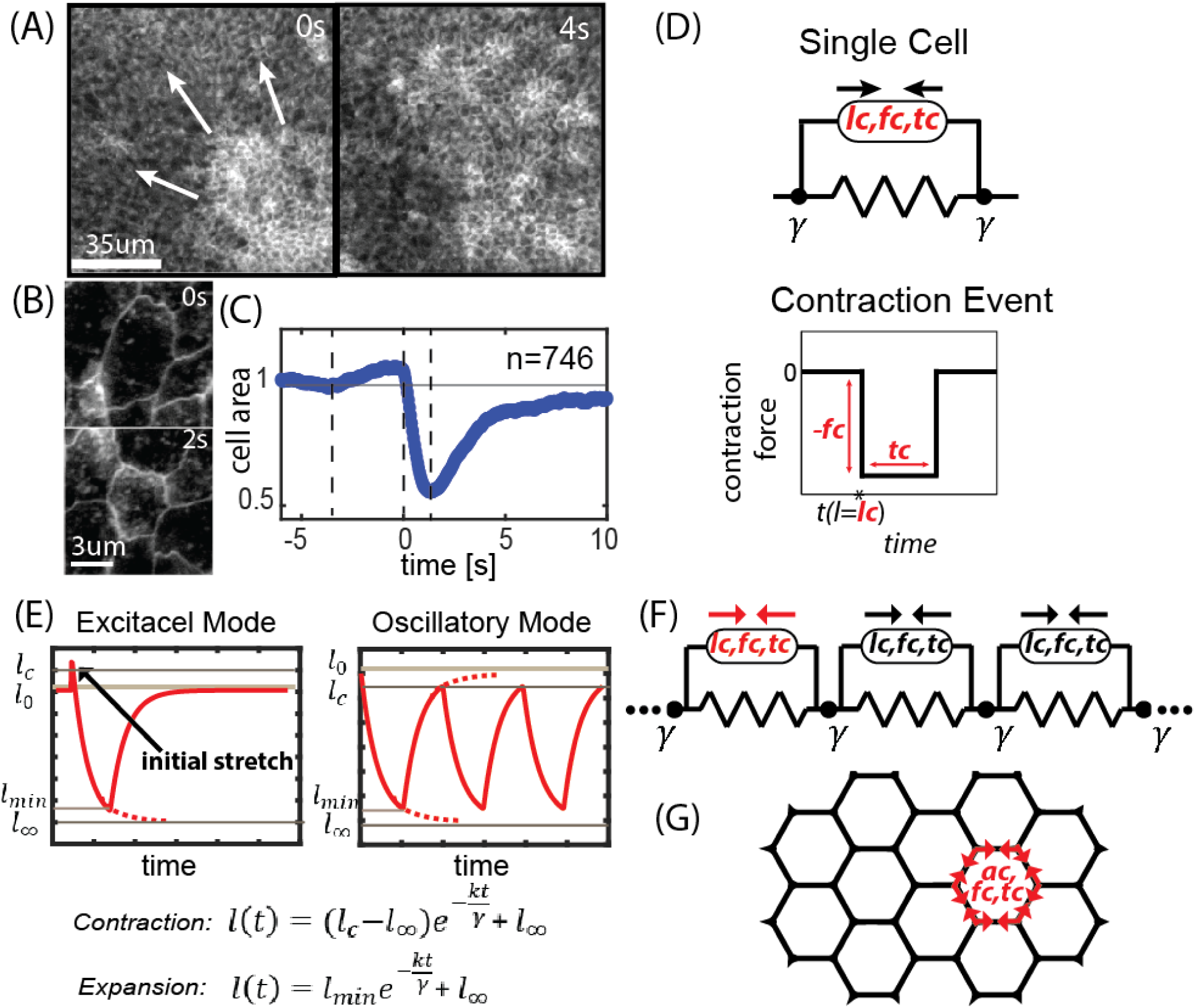
Modeling epithelium with cellular EIC (extension-induced-contraction): (A-C) Experimental results from contractions in the dorsal epithelium of *T. adhaerens* (reproduced from [1]). (A) Snapshots from contraction pulses in the tissue, imaged using fluorescent membrane stain. (B) Snapshots from a cellular contraction event. The contraction induces expansion of neighboring cells. (C) The average contraction profile of single cells within the tissue (such that are not participating in pulses). (D) The mechanical circuit representing the single cell in our model. The active unit, placed in parallel to the spring, represents the active response: beyond a critical length *l*_*c*_, a cell contracts with constant force *f*_*c*_ for a given period of time, *t*_*c*_. In the simulations we use a boxcar function: immediate activation and termination of the force. The cell is found in a media with viscosity *γ* (E) The behavior of isolated such cells, in either an excitable mode (*l*_0_ < *l*_*c*_), or oscillatory mode (*l*_*c*_ < *l*_0_). The piecewise exponential solutions are written below. More details in SI1. (F-G) Sketches of the 1D and 2D settings in our simulations. The degrees of freedom are the vertices between cells that are free to move under cellular forces and viscous drag. The systems are large but finite and the boundary is free.

We now examine the emergent behavior of a finite 1D chain of such cells (fig1f). Each cell is a spring connected in parallel to the active-contractile unit, all experiencing viscous drag from the media. Cell-cell junctions are the nodes, that feel tension due to forces from the nearest cells. Throughout our investigation we choose a finite yet large number of cells, N, and free boundaries, in order to imitate *T. adhaerens*, (details in SI2).

First, we consider a chain of *N* identical excitable cells (i.e. *l*_*c*_ > *l*_0_). Initiating all cells at length *l*_0_, the chain will remain at rest. We now introduce an initial stretch, by initiating a rim-cell at *l*_*i*_ = *l*_*c*_ + *ε*, after which it is set free. The dynamics starts as the stretched cell actively contracts, stretching the next cell in line, due to the spatial coupling in the overdamped conditions. If the induced stretch is sufficient, it triggers another active contraction, and so forth. A contraction pulse then propagates throughout the tissue at a constant speed, *v* (fig2a-b). The emerging longitudinal perturbation is not an acoustic (inertial) wave, but a slower, trigger-wave in an excitable media, that consumes ATP.

The symmetric shape of the strain signal (fig2b-c) may be surprising, as we defined an asymmetric EIC profile for the individual cell: the expansion induces a much larger contraction. In fact, a cell participating in a contraction pulse shows a symmetric strain profile, with a delayed, reduced shrinkage. To understand this emerging effect, let us examine the pulse propagation at the bulk of this large, deeply overdamped tissue. We notice that essentially the only nodes moving are the ones at the interface between active and passive cells - all other nodes are at rest due to force balance. Therefore, at the interface, the sum of the active and passive cell lengths is fixed. Thus, the dynamics in time of a bulk-cell is as follows: (i) starts at *l*_0_ (ii) gets stretched by its active neighbor to exactly *l*_*c*_, and gets activated (iii) contracts until reaching *l*_0_, at which point its passive neighbor reaches exactly *l*_*c*_ and gets activated (iv) our cell keeps applying contraction forces without changing its length, as both its neighbors are also active (v) only when an active neighbor deactivates at *t*_*c*_ and starts to relax - our cell finally shrinks, until reaching *T*_*c*_ itself. As a result, the emergent pulse in the tissue is composed of an extension front, followed by an identical (but opposite-sign) contraction front, and in between, all cells are actively contracting yet remain at their rest-length (fig2a-c, mov1).

Hence, in order to derive a simple scaling for the pulse behavior, as a first approximation, we consider two cells with fixed boundaries (this is practically the case at the interface between active and passive cells). By calculating the time it takes for one cell, contracting with *f*_*c*_, to excite its neighbor 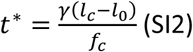, we estimate the propagation speed of the pulse 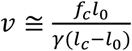, its width *v* ≅ *vt*_*c*_ and its amplitude *amp* ≅ 2(*l*_*c*_ − *l*_0_). Our numerical results confirm all these scaling laws (fig 2h-k).

**Fig 2:**
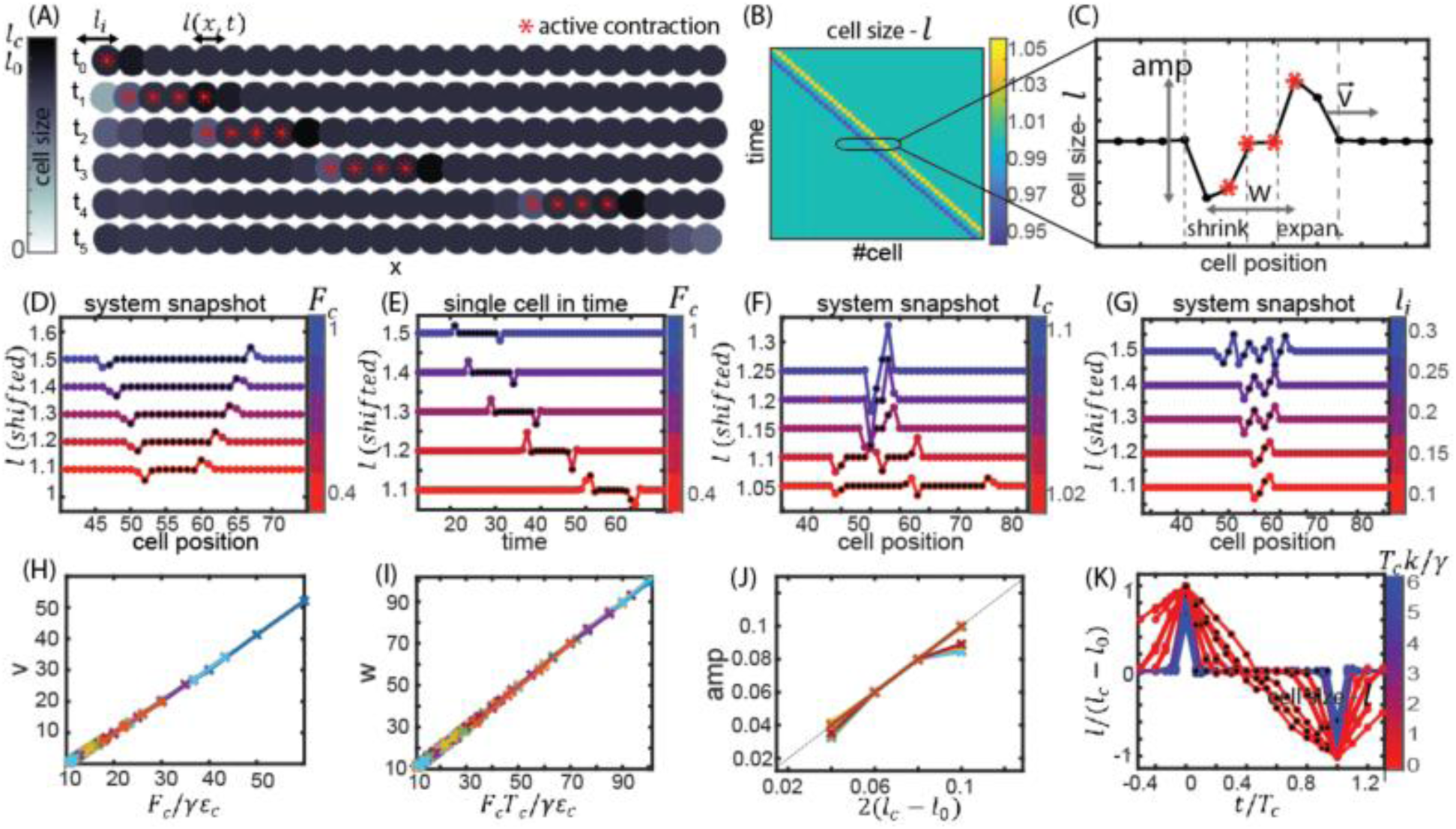
Contraction pulses in a 1D excitable tissue: (A) Snapshots from a dynamic simulation (mov1) of a 1D chain of excitable cells (*l*_0_ < *l*_*c*_) show the propagation of a single contraction pulse from left to right. At *t*_0_ all cells are set at their rest length, *l*_0_, except the first cell on the left, that is initiated at *l*_*i*_ > *l*_*c*_, which triggers the pulse. Greyscale represents cell size; red asters represent activated cells. (B) A kymograph representation of the dynamics shows the pulse propagation in the tissue at constant pulse width, *w*, and speed, *v* (defined in units of cells and cells per time, respectively). (C) A section from the kymograph shows the pulse profile: an expansion-front first, and an equal but opposite contraction-front behind it. All cells in between are actively contracting (marked in red asters) yet found at their rest length. (D) System snapshots from simulations with different *F*_*c*_ values. Results show that Increasing *F*_*c*_ increases the pulse’s width. (E) Time series of a specific bulk cell taken from the same simulations as (D). Results show that increasing *F*_*c*_ increases the pulse’s speed (the pulse reaches the same cell sooner). (F) System snapshots from simulations with different *l*_*c*_ values. Results show that increasing *l*_*c*_ increases the amplitude of the signal. (G) System snapshots from simulations with different *l*_*i*_ values. A “spike train” made of identical pulses is emergent. The number of pulses depends on *l*_*i*_. (H-K) Using the scaling laws we derived, we show a collapse of the numerical results into known functions of the parameters.

We use the derived scaling to find the requirements for pulses to appear: The initial stretch should excite the first cell (*l*_*i*_ > *l*_*c*_), and the impulse of active contraction should be large enough to excite the next cell. Using a set of 3 non-dimensional parameters - normalized time 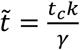, normalized force 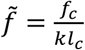, and normalized strain 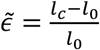 - this requirement estimates to 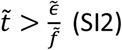. When the system satisfies these criteria, a pulse propagates indefinitely in the tissue with fixed speed. In the absence of these requirements, the initial stretch decays, and bulk cells stay at their rest lengths indefinitely.

Interestingly, we notice that changing the contraction parameters *f*_*c*_, *t*_*c*_ changes the pulse velocity *v* and width *w*, but not the amplitude, *amp* (fig2d-e). Only by changing *l*_*c*_ the amplitude is altered (fig2f). As a result, for a given set of activation parameters (*l*_*c*_, *f*_*c*_, *t*_*c*_), if the external stimulation lasts longer than *t*_*c*_ (in our case, if the initial excitation, *l*_*i*_, is large enough), it generates a “spike train” of several adjacent pulses, all carrying the same quanta of strain - *amp* (fig 2g, SI2). This effect resembles action potential rate encoding behavior in neural networks [33, 34].

Next, we consider a row of N identical oscillatory cells (*l*_*c*_ < *l*_0_). All cells are initiated at *l*_0_ and the boundaries are set to be free. Each cell is a relaxation oscillator, that would beat spontaneously in isolation. However now the cells are additionally subjected to forces coming from their neighbors. We show, that for a wide range of parameters, a mode of ordered pulsations emerges: Initially, all cells contract in a transient irregular phase. Eventually, the system reaches a dynamic steady state (at *T*_*ss*_) where contraction pulses are initiated repeatedly and regularly at the edges and annihilate at the center (fig3a-b, mov1). Annihilation can be seen as a result of the shrinkage front-a “recovering” regime at the back of the pulse that makes the tissue harder to excite (again resembling the refractory period of a neuronal action potential). The annihilation point varies by choosing random initial cell-lengths. The overall tissue length *L*(*t*) shrinks from *L*(*t*_0_) = *N* ∗ *l*_0_ to a steady-state size (*L*_*ss*_) and oscillates around it with amplitude Amp and multiple frequencies *ω*_1,2,3_ (fig 3c).

**Fig3:**
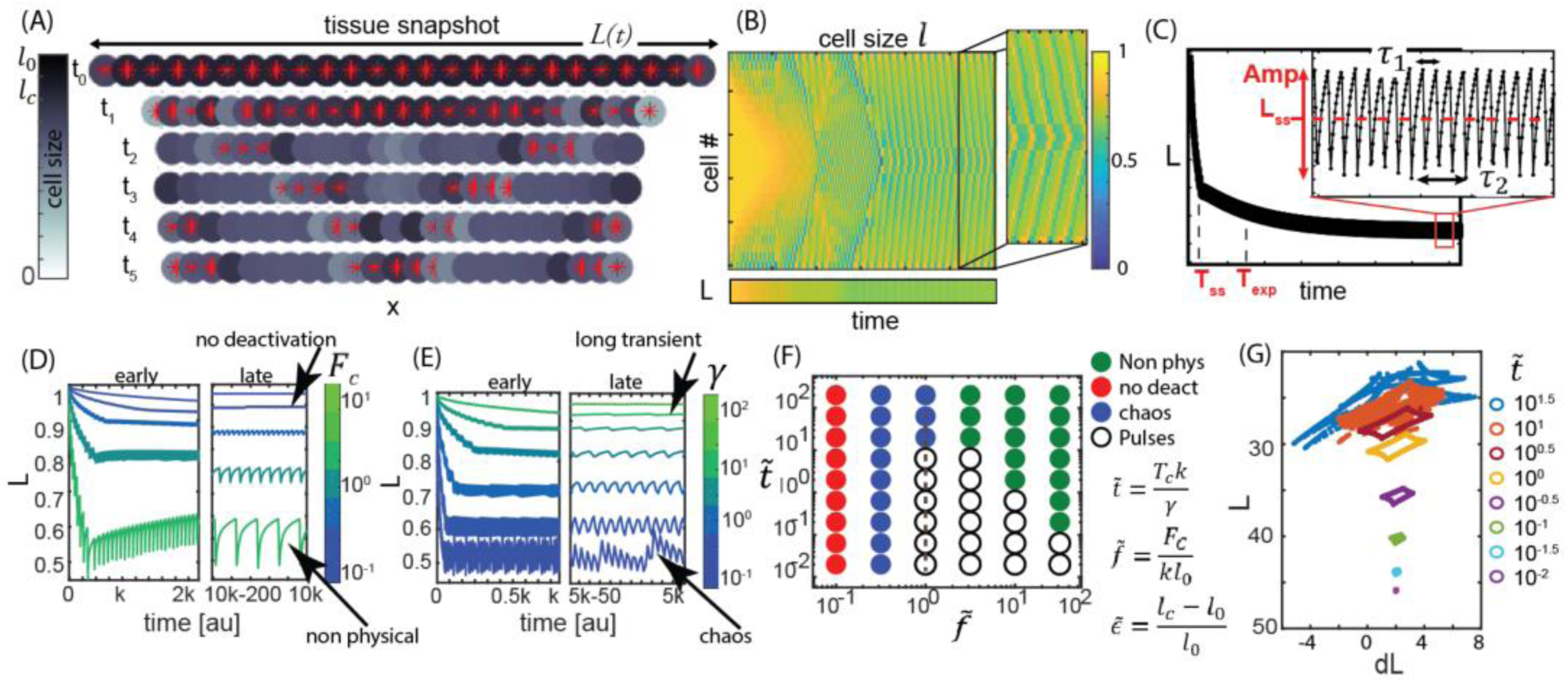
Contraction pulses in a 1D oscillatory tissue: (A) Snapshots from a dynamic simulation (mov1) of a 1D chain of oscillatory cells (*l*_*c*_ < *l*_0_). At *t*_0_ all cells are set to their elastic rest length *l*_0_. At *t*_1_ the system is at its transient shrinking phase. At *t*_2−5_ the system is at its dynamic steady state. Contraction pulses are constantly propagating from the edges to the bulk of the tissue, where they collide and annihilate. The overall length fluctuates around *L*_*ss*_. Greyscale represents cell size; red asters represent activated cells. (B) A kymograph representation of the dynamics shows the pulses propagation at constant speed and their annihilating at the center. The bottom bar shows the overall tissue length, *L*. (C) A sample measurement of *L*(*t*). The ordered traveling pulses emerge at *T*_*ss*_. Then, further exponential decay (time scale *T*_*exp*_) brings the system to its asymptotic average length *L*_*ss*_, around which it fluctuates with multiple frequencies. (D-E) Time series of *L*(*t*) as *F*_*c*_ and *γ* vary. We show early and late intervals, taken from long simulation runs. While we focus on ordered pulsatile behavior, different possible types of dynamics are seen at the extremities of parameter space. These include all-contractile (“no-deactivation”) mode; cell collapse to negative area (“non-physical”); increasingly long *T*_*ss*_ (“long transient”); and irregular traveling pulses (“chaos”). More details in sfig1. (F) A phase diagram shows the different types of dynamics as a function of the non-dimensional parameters 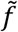 and 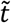. (G) A sharp transition from an ordered pulsation mode (limit cycles) to chaos as a function of *γ*. The trajectory in 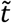 is depicted in dashed line in (F). Plotted data points are taken from steady-state.

In this mode, cells in the bulk (far from both rim and annihilation points) are effectively fixed at a compressed size: due to viscosity and the large number of cells, the time scale for relaxation is much longer than the typical interval between pulses. Therefore, the pulses propagate through a still and uniform background of cells, that are all at *l*∼*l*_*ss*_ < *l*_*c*_. As a result a pulse propagates at a fixed speed. Rim cells are the least constrained and hence relax the fastest after a contraction. When they reach criticality, they initiate a new pulse. The pulse profile features are the same as in the excitable mode. The way the system parameters (*l*_*c*,_*f*_*c*,_*t*_*c*,_*k, γ, N*) relate to the emergent measurables (*L*_*ss*_, *T*_*ss*_, *v, Amp, ω*_1,2,3_) is plotted from our numerical results in fig3d-e and more elaborately in sfig1. Interestingly, most measurables are invariant to the system size (except *T*_*ss*_ and *ω*_2,3_), hence are effective “material parameters”.

Other solutions exist in the oscillatory mode, aside the ordered pulsations, as shown in the phase diagram (fig3f): when the contraction force is weaker than the elastic retraction at *l*_*c*_ (i.e. *f*_*c*_ < *kl*_*c*_) the system will be “stuck”, i.e. continuously apply compression forces but exhibit fixed cell size, that is above *l*_*c*_. These states may look “flickering” with local activation/deactivation close to criticality (mov1). When the active force is too high, we reach a non-physical regime of the model where cells collapse to negative size (in reality, nonlinear elasticity at the limit of compressibility will prevent that). When the viscosity is high, the time it takes to reach steady state is very long, hence in realistic time scales the dynamics may look irregular. Finally, when viscosity is low, irregular pulsations emerge, as a result of various elastic modes that are not overdamped (mov1). The sharp transition between the regular and irregular traveling pulsations is depicted in fig 3g – as a transition from perfect limit cycles to “smeared” chaotic activity.

Finally, we apply the same principles of EIC in a 2D cellular sheet. We use a vertex model, that assumes separate elasticity of a cell perimeter, *k*_*p*_, and a cell area, *k*_*a*_, and is a common modeling approach for confluent tissues. 2D vertex models are controlled by 5 non dimensional parameters [35–37] – a parameter space which we do not examine completely here. However, we do focus on a regime that we believe is most relevant to *T. adhaerens* (low area stiffness *k*_*A*_, high *f*_*c*_, capable of reducing cell area by 50%, and 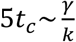). We consider a disk-like shape of a tissue with free boundaries (fig1g). For the single cell rest-shape we choose a regular, compatible hexagon, with rest perimeter *p*_0_ and rest area *a*_0_ (*A*_0_ = 3√3 (*p*_0_/6)^2^/ 2). The cell’s perimeter is controlled by an active unit: when reaching criticality (either a critical area *a*_*c*_ or critical perimeter *p*_*c*_, showing slight differences between them) a compression force *f*_*c*_ is acting for a duration of *t*_*c*_ to shorten the perimeter (see methods).

The results are qualitatively similar to the 1D case: In the excitable mode case (*a*_0_ < *a*_*c*_) a single pulse is propagating from an initially-perturbed point across the tissue (mov2). Note, that the pulse propagates faster closer to the rim. In the oscillatory mode (*a*_*c*_ < *a*_0_), after an initial shrinking phase, the system is self-compressed and contraction pulses are propagating in a uniaxial, azimuthal or a spiral fashion (fig4a, mov3left). As in 1D, an expansion front is a precursor to a shrinkage front, while all cells in between are actively contracting. Static contraction and chaotic modes exist as well, resembling similar states in the 1D case (not shown).

The emergent dynamics we observe take the schematic shape of reaction-diffusion in the displacement vector, *ū*. This can be seen from a simple continuum model written in the spirit of the discrete model we presented here (SI3). The dynamics may be written as: *∂*_*t*_***u*** = ***D***∇^2^***u*** + ***R*** where the diffusion coefficient depend on viscoelasticity |***D***| = *k*/*γ* and the reaction term, ***R***, depends on the cellular activity.

Despite the similarities to the 1D setting, a unique feature of the 2D case is the fact the system is prone to mechanical frustration. An intuitive way to see it, is that rim-cells can only release stresses in the radial axis, but not in the azimuthal one. As a result, rim-cells are not beating like isolated cells, as in 1D. Although rim-cells relax faster than bulk cells, they still need to “wait” for the entire system to relax before they can too, as seen in the radial gradient of strain (mov 3,4,sfig2). In addition, bulk cells are relaxing slowly due to viscosity and due to the energy wells they reach at concave shapes (their perimeter needs to temporarily decrease in order to go back to convexity). The result is long intervals of quiescence. We show that as 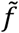 increases, quiescence periods increase, while short bursts of activity occur between them (sfig2).

An intriguing feature of this 2D active tissue is its effective mechanical properties under tensile stress. To demonstrate it, we design a numerical experiment where we pull a 2D sheet at two opposite rim points with a constant force (fig4d, mov4). A passive elastic material at equilibrium focuses high values of stress and strain near the pinching points or along the pulling axis (depending on specifics of the elastic model). The viscous behavior either postpones this elastic equilibrium or creates indefinite creep (again, depending on the specific model used). All these cases impose a threat for tissue cohesion. However, pulling on the active tissue results in a dynamic steady state, where high strains vanish, and instead strains oscillate in time around a lower average, exhibiting both tension and compression (fig 4E). In addition, maximal strains are evenly distributed in space, as all cells are participating in the dynamics (fig4F). This can be seen also in the histogram of all strain values at all times (fig4G). This effect reduces the chances for rupture due to cell strain, at the cost of high values of stress at cell-cell junctions. We bring a full simulation video for the excitable case (*p*_*c*_ > *p*_0_) in mov4.

**Fig4:**
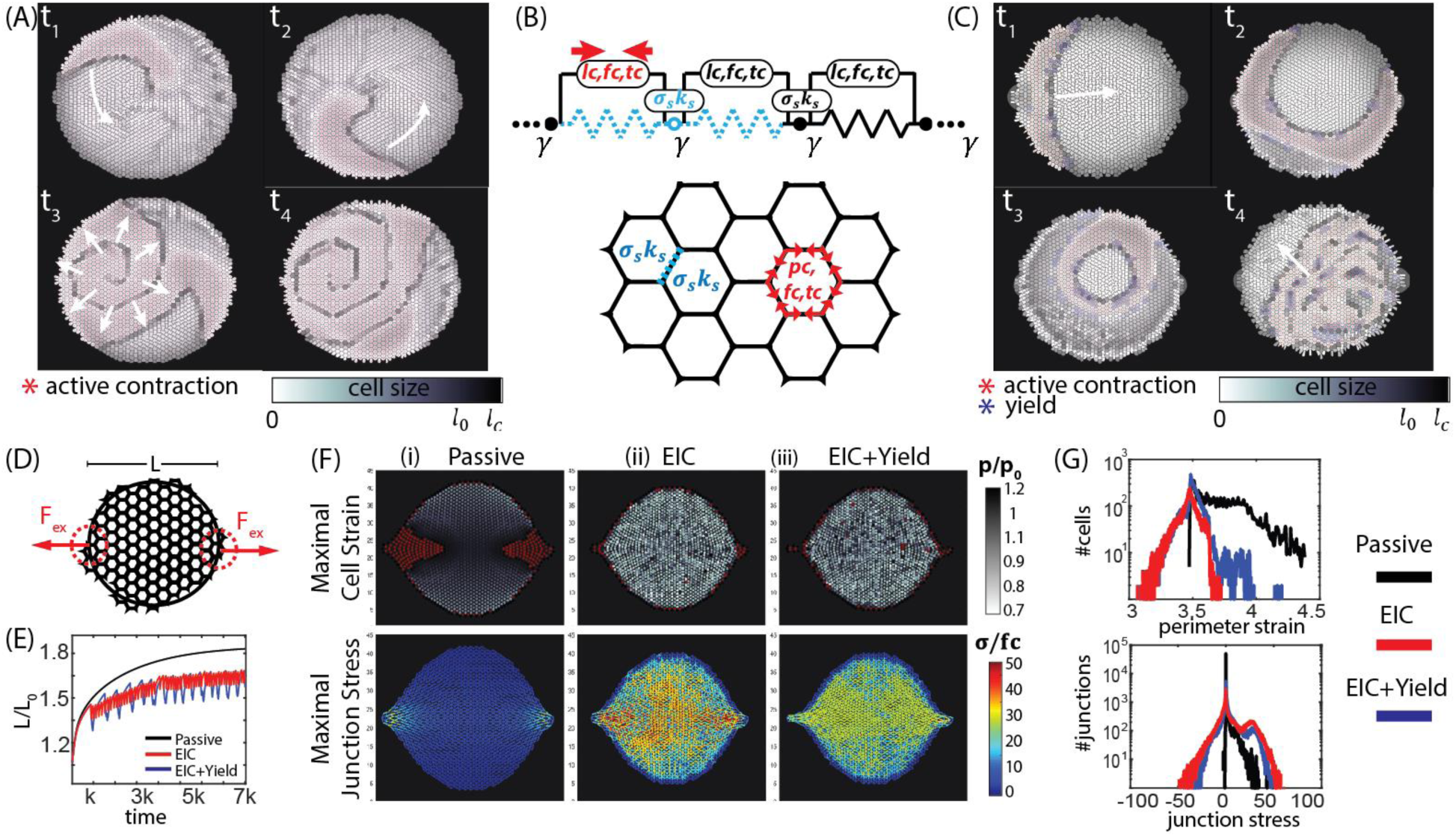
2D Oscillatory tissue: dynamics and “active cohesion”. (A) Snapshots from a dynamic simulation of a 2D cellular sheet with EIC (*a*_*c*_ < *a*_0_, mov 3 left) show circular and spiral pulses with features similar to the 1D case: an extension front, followed by a shrinking front, while all cells in between are actively contracting. Greyscale represents cell area; red asters represent actively contracting cells. (B) Adding a second cellular response: When the tension on a cell-cell junction is higher than a threshold *σ*_*s*_– the cell yields. We model that by immediate softening (*k*_*p*_ → *k*_*s*_) in both neighboring cells. (C) Snapshots from a simulation of a tissue with both contraction and softening thresholds (mov 3 right). The patterns are similar to (A), with softening fronts accompanying the extension and shrinkage fronts. (D-G) Testing the response of an excitable tissue to external stretch in the x-axis: (D) The pulling configuration: constant force is pulling uniaxially on the sheet, acting directly on a fixed set of cells on each side. (E) The distance between the pulling points, L, as a function of time. (F) We compare (i) a passive material (ii) a material with contraction response, (iii) one with both contraction and yielding responses. We present peak values of cell strain (top) and junction stress (bottom) throughout the simulations. The passive material develops strain and stress focusing near the pulling points. The contractile material dynamically distributes the strains in the tissue but “pays” with high values of junction stress. A material with the added yield-response increases slightly the levels of cell strain but cuts-off high junction stress values. (F) Histogram-view of all data points in the pulling simulation.

We show that adding a second cellular response - cell yielding due to junction stress - reduces both maximal junctional stresses and cell strain simultaneously. We add to the simulation a cell softening threshold-decrease in the elastic module *k*_*p*_ by factor two upon reaching critical junction stress *σ*_*s*_ (fig 4B, methods). Once the junction stress is restored below *σ*_*s*_, the stiffness goes back to normal. Results show, that the added response does not change the spatiotemporal patterns significantly, except introducing a tailing softening front (fig4C, mov3). Under external tension, the softening threshold did not change the strain and stress distributions dramatically (mov4), but it did cut-off high stress values, trading them for localized high strains, as seen in the trade-off at the distribution tails (fig 4Eiii,F). When the two thresholds (*a*_*c*_, *σ*_*s*_) are set below rupture values, the tissue is “actively cohesive” as it avoids failure by suppressing both cell strain and junction stress simultaneously.

## Discussion

We suggest a model for propagation of contraction pulses in epithelia. Pulse propagation is a direct result of a single cell’s EIC-contraction due to expansion. Unlike passive elastic retraction after expansion, the model requires the contraction to be active, and include a “memory” time scale, in order to bring a contracting cell significantly below its rest length despite the overdamped conditions. We show that in order for a contraction to propagate in the excitable-cell mode, the contraction impulse should be strong enough relative to the excitation threshold 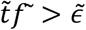 (a qualitatively similar criterion will exist in the oscillatory-cell mode, with *l*_*ss*_ in the role of *l*_0_). The resulting pulses travel in the excitable media via local energy injections, and evolve dynamically following reaction-diffusion equations for the strain.

In actual biological tissues, it is unclear whether an EIC is triggered by strain, stress or strain rate. The exact parameter is hard to distinguish experimentally, and is currently unknown. For our model, we choose to focus on cell-strain and junction-stress as triggers, as they threat tissue cohesion: cell cortex networks rearrange due to stress, but are probable to fail entirely in high strain. Cell-cell junctions, on the other hand, do not “expand”, but may break under stress. In addition, in the presence of drag, a cellular contraction reduces cell strain but increases junction stress. Therefore, we choose high cell strain as the trigger for contraction and high junction stress as the trigger for softening. We claim these feedback loops promote tissue stability.

In this work we choose the active contraction to be governed by a fixed compression force that lasts a fixed duration of time. This is a minimal assumption inspired by our experimental results from *T. adhaerens*, where we showed the single contraction profile, as well as a narrow distribution of contraction times relative to contraction amplitudes and speeds [1]. Nevertheless, other functional behavior of contraction that brings the cell to a new, shrunk size at relevant time scales would create contraction pulses that are qualitatively similar.

We propose a new physiological role for contractility in epithelia: rupture resistance. As EIC is a common epithelial response, and as tissue integrity is at the heart of any epithelium function, “active cohesion” may be relevant in a wide range of systems, even if manifested in different time and length scales. The minimal nonlinear model we presented fits the fast soliton nature of the pulses seen in *T. adhaerens* but may be applicable to other observed contraction waves in epithelia (as in drosophila embryo and MDCK monolayers). This hypothesis should be further tested experimentally. Specifically, it would be interesting to test it in various embryonic tissues, and in epithelia that is either contractile or prone to high, repeatable mechanical stresses (e.g. heart, lung, gut, bladder, vasculature).

Our work brings forward discrete aspects of tissue mechanics, such as cellular chemical thresholds, local mechanical conditions in a contracting cell and preferred cellular geometry. It would be interesting to compare discrete, continuous and hybrid models, to describe observed phenomena in epithelial tissues.

The study of *T. adhaerens* and other early-divergent animals brings tissue-mechanics to the context of evolution of multicellularity. Alongside the fundamental ability of cells to stay cohesive as tissues, it may shed light on the origin of the excitation-contraction coupling and the nervous-muscular system [18]. We suggest to further investigate dynamics in early-epithelia not only in the context of “active cohesion”, but also as possible embodied calculations yielding “behavior” and supporting physiological needs (e.g. locomotion and navigation, wound healing and size control). Finally, this work may inspire engineering of synthetic materials that actively resist rupture.

## Methods

Simulation code was written in MATLAB (MathWorks, 2017b) and is available on GitHub.

We adopt a dynamical modeling paradigm built around gradient-descent on overdamped equations of motion. Unlike conventional use of gradient descent, the energy functional in the algorithm is not constant; rather, it changes every time a cell is activated or deactivated. Inspired by *T. adhaerens*, we take the activation time *t*_*c*_ to be shorter than the viscoelastic time *k*/*γ*, which is the time scale to approach cellular steady state. Therefore, our gradient descent algorithm will not necessarily reach steady state, and indeed in most interesting cases it does not.

The equation of motion is therefore

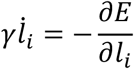

where *l*_*i*_ is the length of cell #i, and the overall free energy *E* is a sum of the energies of all cells, *E*(*t*) = ∑_*i*_ *ε*_*i*_(*t*). (Note that the cellular energy *ε*_*i*_ is different than the cell strain *ϵ*_*i*_ mentioned earlier). In 1D, the energy of each cell is given by

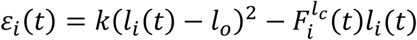

where the boxcar function 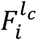 represents the cellular contraction forces: 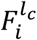 takes the value *f*_*c*_ when *l*_*i*_ > *l*_*c*_ and maintains it for a duration of *t*_*c*_, after which it goes back to 0 (fig1d).

To model 2D tissues, we generalize the above framework using a 2D vertex-model, evolving under the cellular energy function

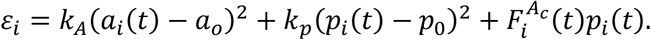

The function 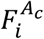 now takes the value *f*_*c*_ when *a*_*i*_ > *a*_*c*_ (or when *p* > *p*_*c*_) and maintains it for a duration of *t*_*c*_. Note that the force may be triggered by an area or perimeter threshold but acts on the perimeter.

We define the stress on cell-cell junction for each edge in the 2D hexagonal grid (an edge is attaching two cells, and is composed of two vertices). We evaluate this stress using a three-step procedure. First, the forces on the two relevant vertices arising from the first cell are averaged, and their mean is projected to the axis normal to the edge. The same average and projection is calculated for forces arising from the other cell. Second, for each pair of cells with a shared edge the two normal forces are subtracted to find the force required to maintain the constraint of confluency. This calculated force is then divided by the edge length to find the force per unit length associated with each junction.

To model tension-induced yielding we add a second cellular response – softening of both cells adjacent to an overstressed junction (we treat cell edges as junctions). The energy functional then becomes:

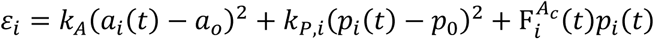

where *k*_*P,i*_ = *k*_*P*_ if the edges of the *i*th cell are all under the critical tension *σ*_*s*_, and *k*_*P,i*_ = *k*_*s*_ < *k*_*P*_ if one of its junctions is overtensed. In the numerical experiments presented here we choose *k*_*s*_ = *k*_*P*_/2. Once the normal stress is reduced below *σ*_*s*_, perimeter stiffness is set back to *k*_*p*_.

## Supporting information

Mov 1

Mov 2

Mov 3

Mov 4

## Contributions

Designed the research: SA, MSB, MP; Wrote the simulations: MSB; Performed the numerical experiments: SA, MSB; Analyzed the data: SA; Wrote the scaling for contraction pulses: HA. Wrote the continuum model: AM; Wrote the manuscript: SA; Supervised the project: MP.

## Acknowledgements

We thank Elisha Moses, Sam Safran, Eran Bouchbinder, Nir Gov and Efi Efrati for useful discussions and comments. We acknowledge financial support from the Weizmann Institute of Science -Women Bridge Position and the Israeli Ministry of Absorption New Immigrant funds to S.A, Horwitz Research grant and the Center for New Scientists at the Weizmann Institute for HA, NIH Innovators award, HHMI Faculty Fellows Program, NSF CCC award and CZI BioHub Investigator funds to M.P.

## Movie Captions

### Mov#1: Different phases in 1D chains

Different dynamic phenomena in 1D chains of cells with EIC. Each circle is a cell. All cells are initiated at their rest length *l*_0_. Cell’s color represents its length: dark cells are longer than white cells. The color scale varies between movies, for clarity. Red asters mark actively contracting cells. Top to bottom: (a) A chain of cells in an excitable mode (*l*_*c*_ = 1.05*l*_0_). We excite the system by initiating the first cell on the left slightly above ℓ_*c*_. The movie is slowed down 50x compared to the rest of the movies (b-e) Chains of cells in an oscillatory mode (*l*_*c*_ = 0.85*l*_0_) exhibit various phenomenology (see fig 3F). (b) Regular pulsations starting from the edges (c) All cells contracting indefinitely, but are “stuck” above *l*_*c*_ (due to low 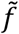) (d) Chaotic behavior close to *l*_*c*_ (due to low 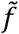) (e) Chaotic behavior due to inertial modes (due to low *γ*).

### Mov#2: 2D excitable sheets

A contraction pulse in a 2D cellular sheet with EIC at the excitable mode (*a*_*c*_ > *a*_0_). We initialize all cells at their rest sizes (*a*_0_, *p*_0_), and excite the system by forcing a single cell to contract (bottom left). As a result, a contraction pulse is propagating in the tissue. The pulse is composed of an expansion front, a shrinkage front, and between them a range where all cells are actively contracting. Face color represents cell area. Red dots mark actively contracting cells.

#### Mov#3: 2D oscillatory sheets

Simulating 2D sheets of cells with EIC at the oscillatory mode. Left: without the softening trigger, Right: with the added softening trigger. Propagating pulses that resemble reaction-diffusion patterns are seen: radial and spiral wave). Also visible are long intervals of quiescence due to mechanical frustration. All cells are initiated at their rest area and perimeter. Face color represents cell area (darker mean smaller cells). Red dots mark actively contracting cells and blue hexagons mark cells that are softened.

### Mov#4: 2D excitable sheets under external tension

Comparing response to tension in 2D sheets of cells. Cell strains are depicted on the left: face color is area-strain, edge color is perimeter-strain. Junction stresses are depicted on the right, each circle represents the normal stress to a cell edge. The external force is applied on the cells in the grey circles and is fixed in time. Top to bottom: (a) A passive sheet reaches equilibrium (b) Tissue with EIC in the excitable mode, using a perimeter-trigger, *p*_*c*_. The tissue is indefinitely dynamic (with area-trigger force balance may be reached). The dynamics include contraction pulses propagating in the direction of the external pull (In the oscillatory mode transverse pulses are more dominant), and periods of slow relaxation (quiescence is more prevalent when *k*_*p*_ > *k*_*A*_). The system is reaching a dynamics steady state, with low cells’ strain but high junctions’ stress. (c) Adding to (b) a yielding response, makes the strain oscillations larger and the quiescence periods longer. As a result, junction stress is lowered compared to (b). All cells are initiated at their rest size. Face color represents cell area, red dots mark actively contracting cells and blue asters mark cells that are softened.

