## Supplementary Information for "Epithelial Tissues as Active Solids: From Nonlinear Contraction Pulses to Rupture Resistance"

### SI 1: Analytics for a single cell

We now formulate the dynamic behavior of a single, isolated cell with an extension-induced-contraction (EIC) response.
We consider a single cell as a spring with rest length $\ell_0$ and stiffness $k$, connected in parallel to an active-contractile unit, characterized by three parameters: $f_c, t_c, \ell_c$. The cell is found in a media with viscosity $\gamma$, in overdamped conditions. The equation of motion for the cell's length, $\ell$, takes the shape:

$$-\gamma\dot{\ell} - k(\ell - \ell_0) = f_{tot} = \begin{cases} f_c & \text{cell is active} \\ 0 & \text{cell is inactive} \end{cases}.$$

A contraction is initiated when $\ell = \ell_c$, either for an excitable cell, i.e. when $\ell_c > \ell_0$, or for an oscillatory cell, i.e. when $\ell_c < \ell_0$. The solution to the above equation is then

$$\ell_{\text{contraction}}(t) = \ell_0 - \frac{f_c}{k} + \left(\ell_c - \ell_0 + \frac{f_c}{k}\right) e^{-\frac{kt}{\gamma}}.$$

In fact, due to the fixed duration in which the contraction is active, $t_c$, the isolated cell does not reach its asymptotic length $\ell_\infty = \ell_0 - \frac{f_c}{k}$, but rather the minimal length

$$\ell_{min} = \ell_{\text{contraction}}(t = t_c) = \ell_0 - \frac{f_c}{k} + \left(\ell_c - \ell_0 + \frac{f_c}{k}\right) e^{-\frac{kt_c}{\gamma}}.$$

Therefore, during relaxation:

$$\ell_{\text{relaxation}}(t) = \ell_0 + (\ell_{min} - \ell_0)\, e^{-\frac{k(t-t_c)}{\gamma}} = \ell_0 + \left(\ell_c - \ell_0 + \frac{f_c}{k}\left[1 - e^{\frac{kt_c}{\gamma}}\right]\right) e^{-\frac{kt}{\gamma}}.$$

In the oscillatory mode, relaxation also lasts a finite period of time, $t_R$, defined by:

$$\ell_{\text{relaxation}}(t = t_c + t_R) = \ell_c,$$

and the frequency of the relaxation-oscillator is then:

$$f_1 = \frac{1}{t_R + t_c}.$$

### SI 2: Analytics for a 1D excitable system

We now formulate the properties of a single pulse propagating in a 1D excitable system. We consider a row of N cells (fig 1f in the main text) in an excitable mode (i.e. with $\ell_c > \ell_0$). Cell-cell junctions are the vertices. We assume that the drag from the media acts on all vertices directly (no distinguished intra-cellular viscosity). We have in the problem 6 parameters:

$$\ell_c, f_c, t_c, \gamma, k, \ell_0$$

and 3 unit-types: force, length and time, hence 3 non-dimensional groups. We choose the set:

$$\tilde{\epsilon} = \frac{\ell_c - \ell_0}{\ell_0} \;;\; \tilde{f} = \frac{f_c}{k\ell_0} \;;\; \tilde{t} = \frac{t_c k}{\gamma}$$

**Model assumptions, taken from Tplax**:

1. Contraction threshold is not far from the rest length:

$$\ell_c - \ell_0 < 1$$

2. Activity is stronger than elasticity at contraction initiation:

$$F_c \gg k(\ell_c - \ell_0)$$

3. Activity is not strong enough for the cell's length to become negative:

$$\ell_{min} \geq 0$$

**Therefore, for the excitable-cell mode, we ran simulations in the following regimes**: $\ell_0 = 1; k = 1 : 2; f_c = 0.25 : 2$ (once too low - there will be no propagation, once too high - cells collapse. For simplicity we keep $\ell_\infty > 0$); $\ell_c = 1.02 : 1.05$; $t_c = 1.8 : 3$ (if too high - $w > N$, if too low - $w < 1$); $\gamma = 2 : 20$ (if too high - $t_r > t_c$, if too low - we deviate from the overdamping assumption).

#### Criterion for Propagation and Propagation Speed

In order to estimate the pulse propagation speed, let us examine the propagation at the bulk of a large, deeply overdamped tissue. We notice that essentially the only nodes moving are the ones at the interface between active and passive cells - all other nodes are at rest due to force balance. Therefore, at the interface, the sum of the active and passive cell lengths is fixed (for a more complete solution, one should solve for the object that is indefinitely long chain of springs- one on each side). Taking that estimation, we consider a tissue of two cells at rest, fixed at the boundary. When cell#1 actively contract, the other one expands accordingly.

Let $x$ denote the displacement of the interface vertex. At $t = 0$, cell #1 is activated as its length hits $\ell_c$, and cell #2 is still at its rest length $\ell_0$. The force acting on $x$, composed of an active force from one side and two restoring spring forces, then generates

$$\dot{x} = \frac{f_{tot}}{\gamma} = \frac{f_c - 2kx}{\gamma},$$

whose solution is

$$x = \frac{f_c}{2k}\left(1 - e^{\frac{-2kt}{\gamma}}\right).$$

Cell #2 will be activated when

$$\ell_c - \ell_0 = x^* = \frac{f_c}{2k}\left(1 - e^{\frac{-2kt^*}{\gamma}}\right),$$

thus at time

$$t^* = -\frac{\gamma}{2k}\log(1 - \frac{2k(\ell_c - \ell_0)}{f_c}),$$

if the activation time of cell #1 did not end yet by that time. Thus, the criterion for pulse propagation is

$$t_c > -\frac{\gamma}{2k}\log(1 - \frac{2k(\ell_c - \ell_0)}{f_c}),$$

or in dimensionless form

$$\tilde{t} > -\frac{1}{2}\log(1 - \frac{2\tilde{\epsilon}}{\tilde{f}}).$$

Under assumption (2), and using the Taylor expansion $\log(1+x) \approx x$ for $x \ll 1$, the propagation criterion becomes

$$\tilde{t} > \tilde{\epsilon}/\tilde{f}.$$

The activation of cell #2 occurs at

$$t^* \approx \frac{\gamma(\ell_c - \ell_0)}{F_c}$$

and the pulse speed through the bulk will then be

$$v = \frac{\ell_0}{t^*} \approx \frac{f_c \ell_0}{\gamma(\ell_c - \ell_0)} = \frac{\ell_0 k}{\gamma}\frac{\tilde{f}}{\tilde{\epsilon}}$$

#### More emerged quantities

The pulse width is the distance in space between the expansion peak and the contraction peak. It is also the distance the pulse propagates at time $t_c$, thus the length of the consecutive active tissue

$$w = vt_c$$

The amplitude of the pulse is the difference between the maximal and minimal cell lengths, or the amount of strain the pulse carries:

$$amp \approx 2(\ell_c - \ell_0)$$

(When an active cell reaches $\ell_0$, it activates its passive neighbor and then stops contracting due to force balance).
The number of sequential pulses in a "spike train" is the ratio between the duration of the external stimulation, $t_A$ and the activation time, $t_c$. In our case, we apply this stimulation by setting the boundary cell to an initial length $\ell_i$. Then, $t_A$ is the time it takes this cell to contract back to below $\ell_c$, therefore

$$\#init = t_A/t_c = (\frac{\gamma}{k}\log\frac{\ell_i - \ell_\infty}{\ell_c - \ell_\infty})/t_c.$$

#### SI 3 : A continuum model

In this short supplementary information section, we propose a continuum model analogous to the discrete model used in the main text. The suggested model first invokes momentum balance as

$$\rho\ddot{\boldsymbol{u}}(\boldsymbol{r},t) + \eta\dot{\boldsymbol{u}}(\boldsymbol{r},t) = \nabla\cdot\boldsymbol{\sigma}(\boldsymbol{r},t),$$

where $\boldsymbol{u}(\boldsymbol{r},t)$ is the displacement vector, $\rho$ is the mass density, $\eta$ is the viscosity, and $\boldsymbol{\sigma}(\boldsymbol{r},t)$ is the Cauchy stress tensor (we suppress the explicit time and spatial dependence $(\boldsymbol{r},t)$ from here on).

As the system is assumed to be over-damped, we neglect the inertial term $\rho\ddot{\boldsymbol{u}}(\boldsymbol{r},t)$, and the above equation reduces to

$$\eta\dot{\boldsymbol{u}} = \nabla\cdot\boldsymbol{\sigma}\,.$$

We now need to specify a constitutive relation between the Cauchy stress tensor $\boldsymbol{\sigma}$ and the displacement $\boldsymbol{u}$ to describe material response. Inspired by the discrete model presented in the main text, we decompose the stress tensor $\boldsymbol{\sigma}$ in accordance to the Kevlin-Voigt mode, as

$$\boldsymbol{\sigma} = \boldsymbol{\sigma}^{el} + \boldsymbol{\sigma}^{ac}\,,$$

where $\boldsymbol{\sigma}^{el}$ is the elastic component, and $\boldsymbol{\sigma}^{ac}$ is the active one. We take the elastic stress to depend linearly on the strain $\boldsymbol{\epsilon} \equiv \frac{1}{2}(\nabla\boldsymbol{u} + (\nabla\boldsymbol{u})^T)$, generally written as $\boldsymbol{\sigma}^{el} = \boldsymbol{C}\nabla\boldsymbol{u}$ (here $\boldsymbol{C}$ is stiffness tensor). Currently we leave the active stress $\boldsymbol{\sigma}^{ac}$ unspecified, though it is clear that it may depend on the strain $\boldsymbol{\epsilon}$ as well as additional internal cellular properties (e.g. chemical agents, protein concentration etc.) denoted here as $\mathcal{I}_\alpha$ (where $\alpha$ enumerates the different properties used), $\boldsymbol{\sigma}^{ac}(\boldsymbol{\epsilon},\mathcal{I}_\alpha)$.

Using the stress decomposition in the momentum balance equation yields a dynamic equation for the displacement $\boldsymbol{u}$ as

$$\dot{\boldsymbol{u}} = \nabla\cdot\boldsymbol{D}\nabla\boldsymbol{u} + \boldsymbol{R}(\boldsymbol{\epsilon},\mathcal{I}_\alpha)\ \,,$$

where $\boldsymbol{D} \equiv \boldsymbol{C}/\eta$ is an effective diffusion tensor, and $\boldsymbol{R}(\boldsymbol{\epsilon},\mathcal{I}_\alpha) \equiv \frac{1}{\eta}\nabla\cdot\boldsymbol{\sigma}^{ac}(\boldsymbol{\epsilon},\mathcal{I}_\alpha)$ is the reaction term.

For simplicity, from this point on we consider the 1D, scalar analogue of the above equation, written as

$$\dot{u} = D\partial_{xx}u + R(\epsilon,\mathcal{I}_\alpha)\ \,,$$

where $u$ is a scalar field, $\epsilon \equiv \partial_x u$, $R(\epsilon,\mathcal{I}_\alpha) \equiv \frac{1}{\eta}\partial_x\sigma^{ac}(\epsilon,\mathcal{I}_\alpha)$ is the scalar analogue of $\boldsymbol{R}(\boldsymbol{\epsilon},\mathcal{I}_\alpha)$, and $D$ is a diffusion coefficient. The active reaction term $R(\epsilon,\mathcal{I}_\alpha)$ should be supplemented with an evolution equation for the internal cellular properties $\mathcal{I}_\alpha$. While there are numerous ways to specify the dynamics of $\mathcal{I}_\alpha$, we focus on the following generic equation of motion

$$\dot{\mathcal{I}}_\alpha = \frac{1}{\tau_\alpha}g_\alpha(\boldsymbol{\epsilon},\mathcal{I}_\alpha)\,,$$

where $\tau_\alpha$ is a typical time-scale for the dynamics, and $g_\alpha(\boldsymbol{\epsilon},\mathcal{I}_\alpha)$ captures the dynamics of $\mathcal{I}_\alpha$ properly. In the proposition above cell-cell coupling is purely mechanical, i.e. the functions $g_\alpha$ do not include spatial derivatives of any sort.

We now consider the proposed equations with a single cellular property $\mathcal{J}$ as it is sufficient in order to capture the phenomenology of the discrete model proposed in the text. As a consequence, we introduce only a single time-scale $\tau$, which can be mapped to the force-duration time $T_C$ in our discrete model.

Next, we simplify reaction term $\sigma^{ac}$ by assuming it depends on single the cellular component $\mathcal{J}$ only. Explicitly, we assume it to have the form

$$\sigma^{ac}(\mathcal{J}) = \begin{cases} 0, & \mathcal{J} < \mathcal{J}^* \\ -f, & x \geq \mathcal{J}^* \end{cases},$$

where the stress' amplitude $f$ is analogous to $F_C$ in the discrete model, and $\mathcal{J}^*$ is a sharp activation cut-off.

Finally to relate between the sharp cut-off $\mathcal{J}^*$ and the critical length $L_C$ in the discrete model, we use $g(\epsilon, \mathcal{J})$ to retrieve a dimensional factor $\kappa$ with similar dimensions to $\mathcal{J}$, such that $\epsilon^* \equiv \frac{\mathcal{J}^*}{\kappa}$. By considering a single cell property $\mathcal{J}$, we have shown the dynamical equations above supply similar parameters as those used in our discrete model, and that the two could be cast on an equivalent footing.
